# FluxNorm: Toolbox for metabolic flux assay normalization by *in situ* cell counting

**DOI:** 10.1101/2023.10.13.562314

**Authors:** Nathalie A. Djaja, Teva Bracha, Seungyoon B. Yu, Haoming Wang, Natasha M. Carlson, Gulcin Pekkurnaz

**Author notes:** These authors contributed equally to this work (Nathalie A. Djaja, Teva Bracha).

## Abstract

Plate-based quantitative metabolic flux analysis has emerged as the central technology to examine cellular metabolism and mitochondrial bioenergetics. However, accurate interpretation of metabolic activity between different experimental conditions in multi-well microplates requires data normalization based on *in situ* cell counts. Here, we describe FluxNorm, a platform-independent semi-automated computational workflow, validated for three different cell types, to normalize cell density for accurate assessment of cellular bioenergetics.

---

Metabolic flux analysis using Agilent’s Seahorse Bioanalyzer has emerged as a crucial tool for evaluating cellular metabolism and mitochondrial bioenergetics. Despite its prowess, challenges persist in normalizing metabolic flux data to achieve precise results. The XF metabolic flux analyzer is a high-throughput plate-based assay that measures oxygen and proton flux to calculate oxygen consumption rate (OCR) and extracellular acidification rate (ECAR), respectively. However, cell density often varies between wells and causes discrepancies in absolute measurements. Assuming a consistent plating density across experimental conditions is a fundamental flaw as certain manipulations can inherently affect cell density (i.e. by changing the cell proliferation rate). Currently, three methods are commonly used for normalizing seahorse metabolic flux data: protein measurement, genomic DNA measurement, and cell counting ^1-4^. Both protein measurement and genomic DNA measurement require biochemical assays following metabolic flux experiments^5^, and may introduce sample transfer error or be incompatible with certain experimental manipulations. For example, protein measurements cannot be used if mechanisms involving protein synthesis are targeted – these are often used to investigate mitochondrial biogenesis^6^. Cell counting overcomes the weaknesses of protein or genomic DNA normalization; however, there are practical challenges due to the inaccessibility of automated instruments and the excessive time needed to count cells manually.

Semi-automated cell counting methodologies to obtain quick and accurate cell counts without the need for automated instrumentation are advancing^7,8^. To apply these approaches towards normalization of metabolic flux data, we developed FluxNorm, which is accessible, free, and easy to use. FluxNorm consists of two smaller programs based entirely in freeware: NucJ, an ImageJ macro which counts cells from fluorescent images, and ExtraPy, a short python script which averages and extrapolates the counts from NucJ to represent the entire well (Fig. 1) (https://github.com/pekkurnazlab/FluxNormalyzer). Both programs require calibration and configuration for different cell types. Once the starting parameters for a given set of experimental conditions are established, a trained user can go from having unprocessed fluorescent images, acquired by an inexpensive epifluorescence microscope, to normalized metabolic flux data in several minutes. FluxNorm is composed entirely of freeware, making it an accessible and consistent means to generate cell counts for various cell types including primary neuron cultures (Fig 2).

**Figure 1.**
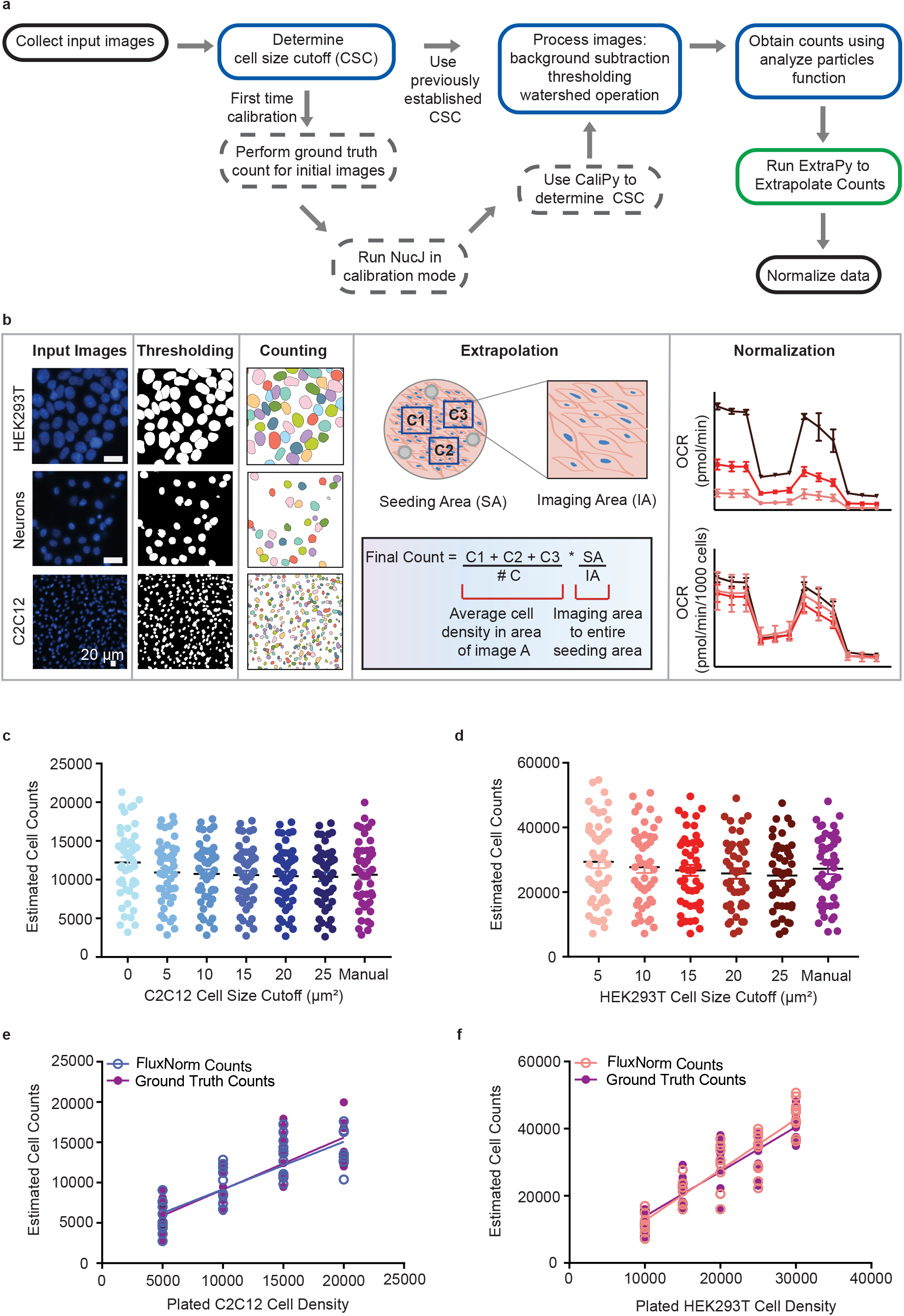
FluxNorm workflow and process. **a**, The FluxNorm pipeline for data normalization. Input images from multi-well plates are manually captured using an epifluorescent microscope. For each novel cell line, a one-time CSC determination is necessary for optimal cell counts. Subsequently, metabolic flux data is normalized using these cell counts. **b**, Sample input images demonstrating nuclei-stained HEK293T cells, primary cortical neurons, and C2C12 cells (first column) are processed through thresholding and background subtraction (second column) to derive cell counts per image (third column). Densities calculated from sampled images are extrapolated to provide a quantification of the total number of cells per well (fourth column). Seahorse metabolic flux assay measurements, represented as Oxygen Consumption Rate (OCR), are displayed both before normalization (top graph, last column) and post-normalization (bottom graph, last column). **c**,**d**, Cell counts obtained using increasing CSCs for C2C12 (c) and HEK293T (d) cells are plotted and compared to manually obtained ground truth counts (purple) (n= 3 independent experiments, 8 wells per condition). **e, f**, Upon determining the optimal CSC, cell counts at varying plated cell densities for C2C12 and HEK293T cells, as determined by FluxNorm, are plotted against manually counted ground truths (purple). Data are presented as mean ± SEM, (e) p = 0.995 and p = 0.709

**Figure 2.**
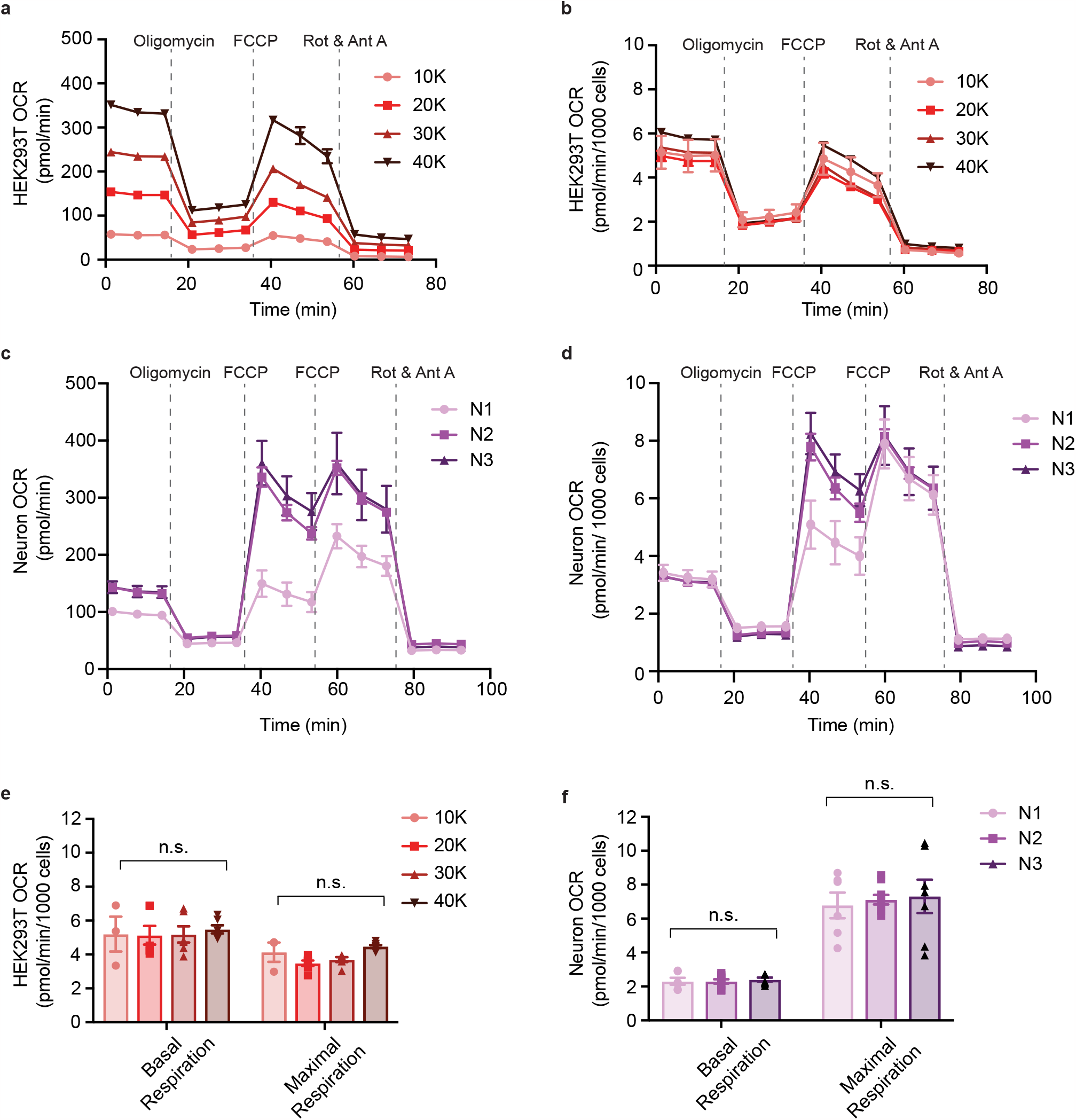
Normalization of metabolic flux data using FluxNorm. **a**,**b**, HEK293T cells exhibit density dependent OCR measurements following the Mito Stress Test protocol (a). This variability is reconciled through FluxNorm-based cell count normalization (b). **c**,**d**, Primary cortical neuron cultures, obtained from three independent preparations but plated at identical densities, exhibit varied OCR readings (c). Following cell count normalization, these cultures display similar OCR values (d). **e**,**f**, Post-FluxNorm normalization, basal and maximal OCR readings remain consistent across varying HEK293T cell densities and three replicate primary neuron preparations. Data are presented as mean ± SEM; p > 0.05, two-way ANOVA.

Metabolic flux assays are a plate-based assay commonly run on 24- or 96-well plates. Due to inconsistencies in plating density that may occur between wells, our methodology involves staining and counting cells to normalize flux data for adherent cell monolayer. Staining cells for cell counting after metabolic flux assay provides its own set of complications. We found that when we fixed, stained, and counted cells upon completion of the assay, cell density often decreased due to excessive perturbations to the cells from washing and removing media. In addition, cells were especially sensitive following the metabolic flux assay because injection of drugs such as rotenone and antimycin A is toxic to cells over time. To overcome this challenge, we added nuclei dye (NucBlue) to the wells before starting the metabolic flux assay. This method avoids perturbations and enables immediate imaging of cells after the assay. We found no difference in metabolic activity when incubating cells with NucBlue prior to the assay (Supp Fig 1), and enhanced staining with reduced background due to longer incubation time.

Counting cells on Agilent’s microplates presents unique challenges compared to imaging regular microwell plates due to the presence of three “standoffs” within each well. While these standoffs allow sensors to capture metabolic flux measurements, they must be avoided during image acquisition. Cells tend to clump around these standoffs, and the fluorometric sensor on the cartridge does not cover this area for measurements^9^. Rather than counting the entire well while avoiding the standoffs, we sampled representative areas of each well to extrapolate cell counts for the entire well (Fig 1B). This methodology is used by automated instruments and accurately captures cell density^10^. Images used for extrapolation should not contain standoffs or have excessive cell clumping or vacancy. The collection of three representative images per well takes less than an hour for a 96-well plate, making acquisition of cells images significantly faster than dedicated automated cell counting instruments which often take hours.

Reliable normalization of metabolic flux data by FluxNorm requires (1) cell counts generated by NucJ are accurate and (2) differences in metabolic flux data directly relate to plating density. To ensure that NucJ could generate accurate semi-automated cell counts, we implemented a user-defined cell size cutoff (CSC) which excludes particles below a minimum area. This parameter is tuned for cell type and imaging condition via a first-time calibration step (Fig 1A). Briefly, calibration entails manually counting cell nuclei from acquired images to generate ground truth cell counts, then comparing those to counts obtained by NucJ at a range of CSCs (Supp Fig 2). We performed this calibration in C2C12 and HEK293T cells (Fig 1C-F, Supp Fig 3). C2C12 and HEK293T cells were plated at a range of densities, ranging from 5K to 40K cells per well. We then acquired a set of images from these wells and manually counted cells. These manual counts were then compared with counts obtained at a range of CSCs. The CSC which produced the least significant difference from our manual counts was used (Fig 1E-F). This critical step ensures NucJ’s accuracy by comparing automated cell counts with ground truth data.

We next tested FluxNorm’s ability to accurately normalize metabolic flux data. To do this, we plated HEK293T cells at a range of densities and performed Agilent’s Seahorse XF Mito Stress Test. As expected, unnormalized OCR values correlated directly with plating density (Fig 2A). Upon normalization with FluxNorm, OCR values were not statistically different among differing plating densities (Fig 2B, E). We also performed metabolic flux assays and analysis with FluxNorm on primary cortical neuron cultures. Neurons were isolated and plated from three different dissections, resulting in three experimental conditions. Unlike HEK293T cells which can be plated at varying cell densities, neuronal cultures are more sensitive to changes in seeding density. Therefore, we assayed neurons from separate preparations to highlight natural variability between experimental replicates that leads to differing metabolic flux data. Neurons had differing OCR values (Fig 2C) corresponding to varied cell counts prior to normalization. Upon normalization, however, OCR values were comparable among the three differing neuron cultures (Fig 2D, F). Thus, FluxNorm can normalize metabolic flux data from multiple cell types and across experimental conditions to compare oxygen consumption rates when cell density is inconsistent.

The field of metabolic flux analysis is rapidly developing with immense applications for both basic and biomedical research^11,12^. However, a widely accessible and standardized methodology for data normalization has not been established. FluxNorm addresses the shortcomings found in previous cell counting methodologies by providing accessible freeware that is accurate and efficient. We propose FluxNorm as a tool to fill this gap in the field and provide a consistent methodology to normalize metabolic flux data.

## FIGURE LEGENDS

**Extended Data Figure 1.**
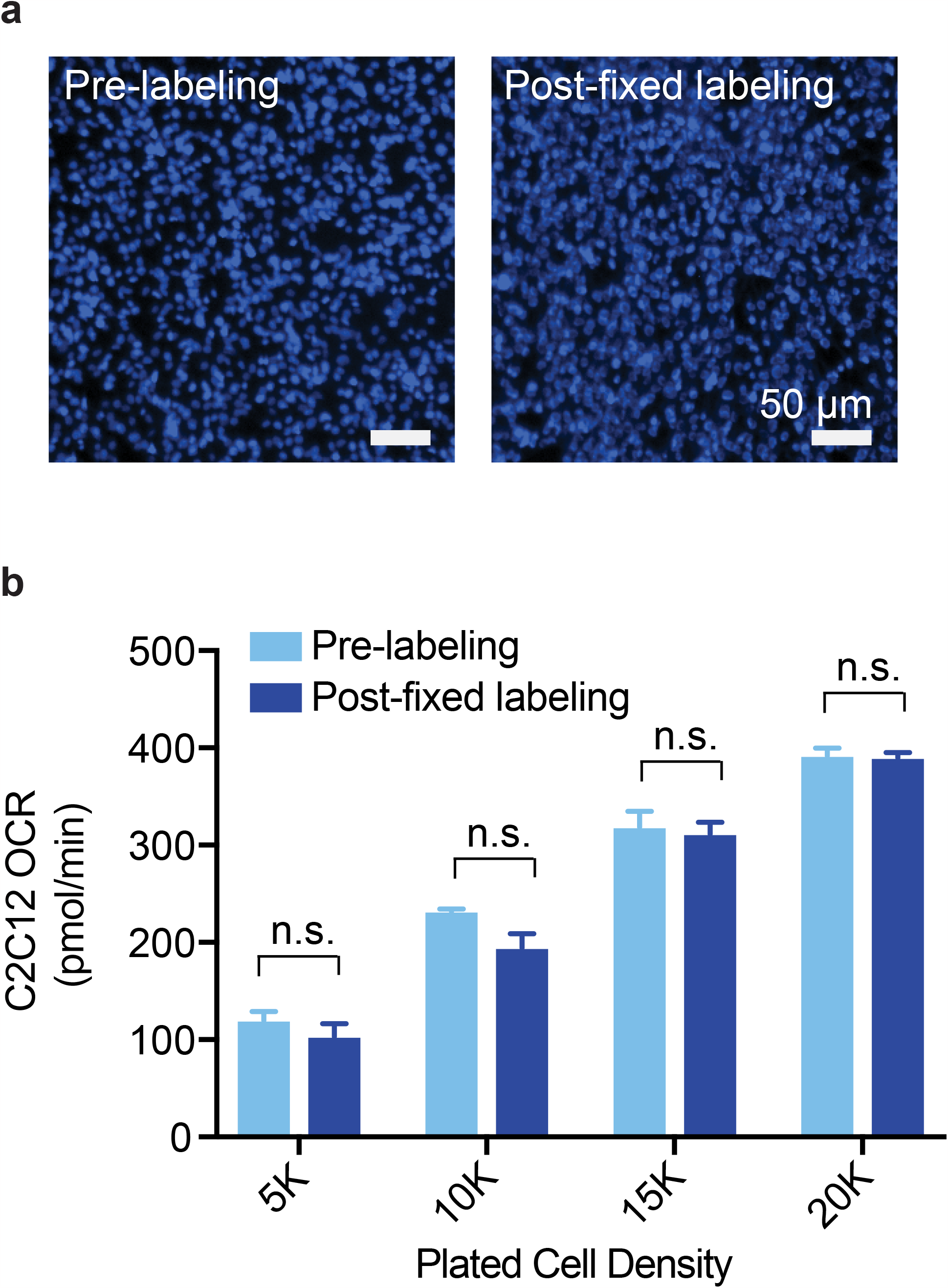
Comparison of live and fixed nuclei labeling for normalization. **a**, C2C12 cells were either pre-labeled with NucBlue prior to the assay or post-labeled after assay completion (post-fixed labeling). Scale bar= 50 μm. **b**, Data normalization using both pre- and post-fixed labeling methods demonstrates no significant variations in metabolic measurements across different C2C12 cell densities. Data are presented as mean ± SEM; p > 0.05, one-way ANOVA.

**Extended Data Figure 2.**
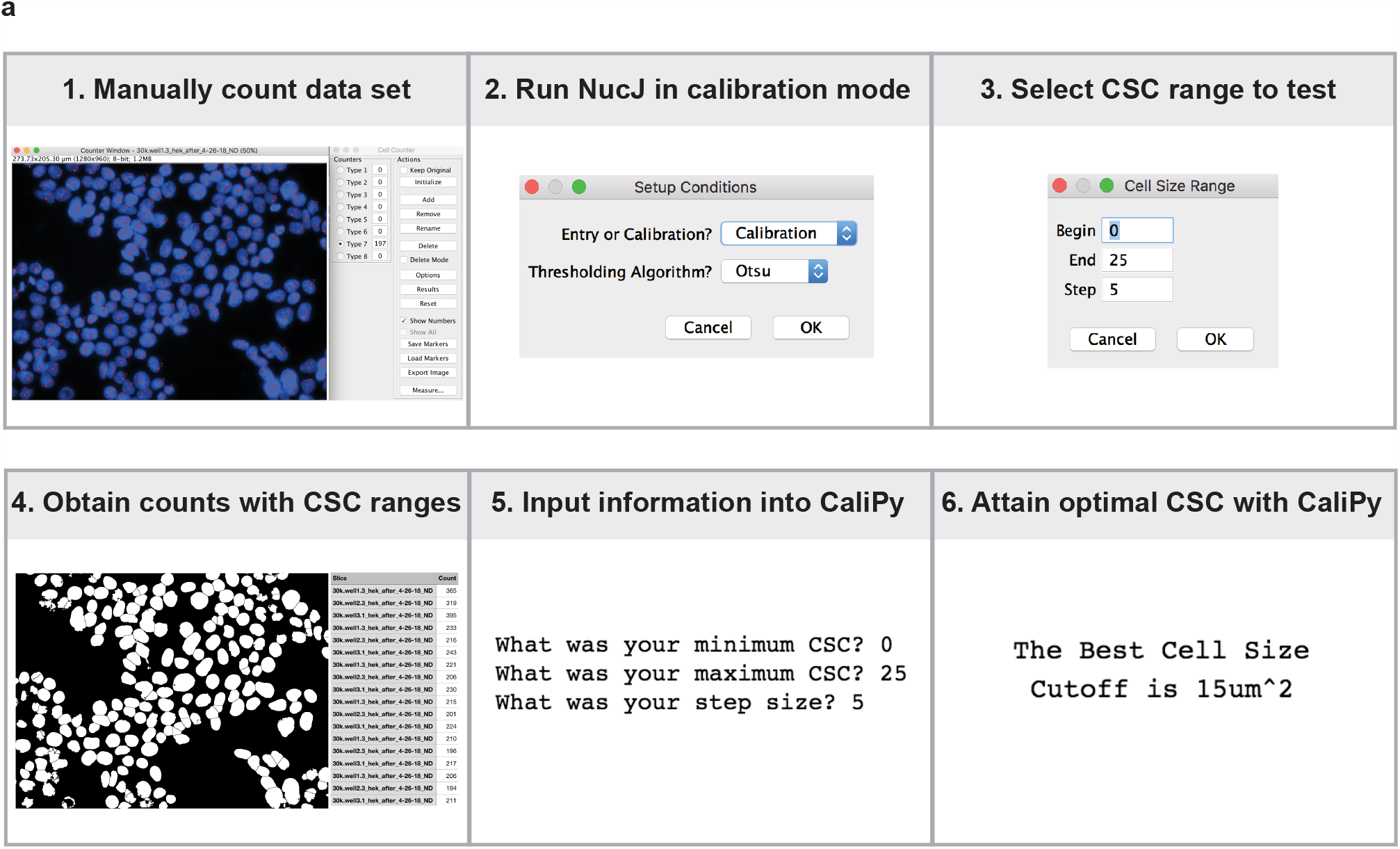
Determination of Optimal Cell Size Cut-off (CSC) for Novel Cell Types Using CaliPy. **a**, Step by step overview of CaliPy. Data sets are first manually counted by the user to obtain an accurate cell count for each input image. NucJ was run in calibration mode. A range of CSCs were chosen with a desired step size. Cell counts will be generated with the range of CSCs selected. CaliPy will then take cell counts generated with varying CSCs and compare to manually obtained GroundTruth counts to determine the optimal CSC.

**Extended Data Figure 3.**
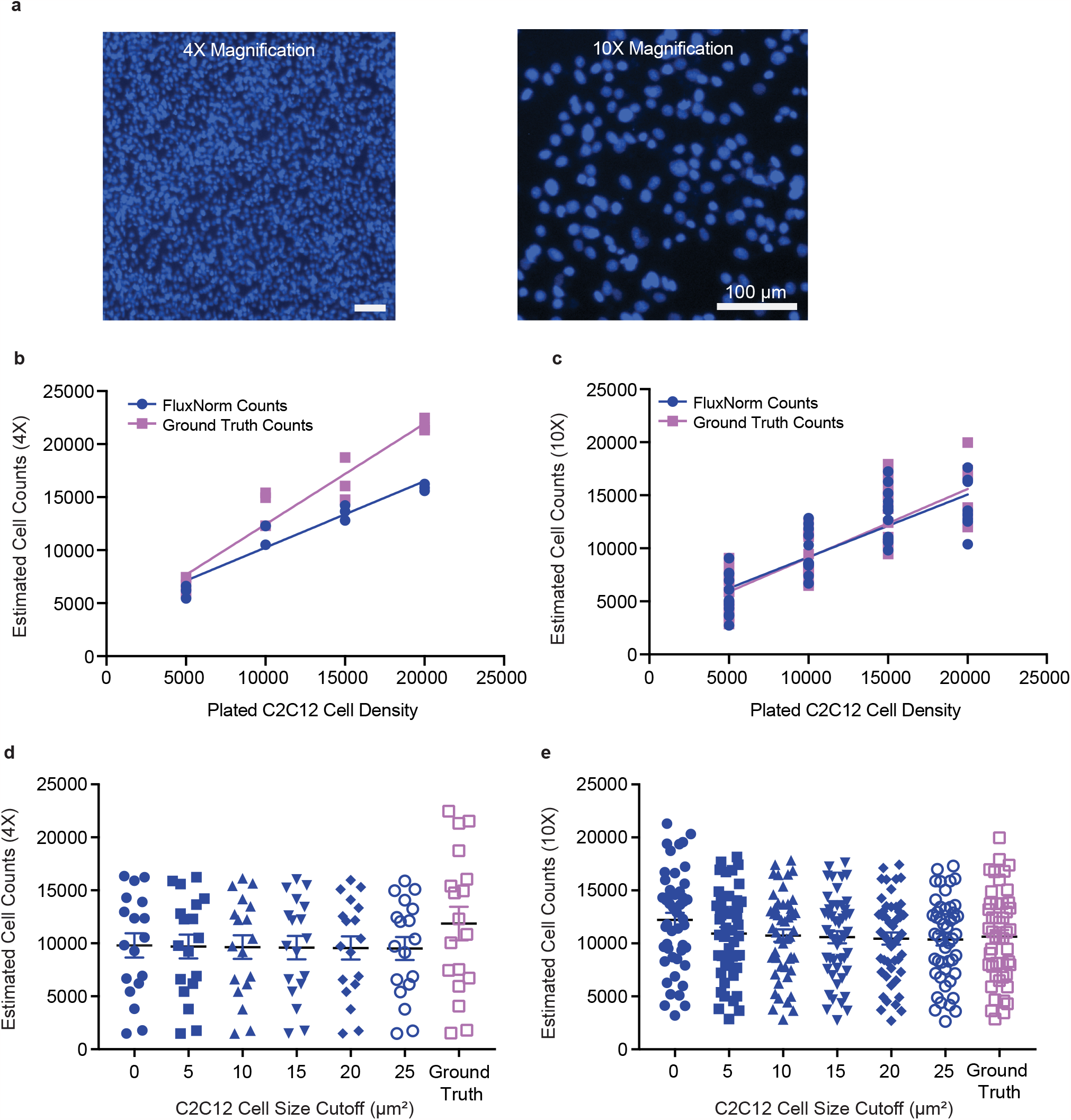
Determination of Optimal Image Acquisition Parameters for FluxNorm. **a**. C2C12 cells captured at both 4X and 10X magnification. Scale bar= 100 μm. **b**,**c**, Comparison of C2C12 cell counts obtained through FluxNorm (blue) with manual GroundTruth counts (purple) from images acquired at 4X (b) and 10X (c) magnification. **d**,**e**, For 4X images, FluxNorm counts fail to align with GroundTruth counts at any chosen CSC (d). However, with enhanced resolution at 10X magnification, FluxNorm counts closely match the GroundTruth counts (e). Data are presented as mean ± SEM.

## METHODS

### Animals

All animal housing, breeding, and procedures were conducted according to the NIH Guide for the Care and Use of Experimental Animals and approved by the University of California San Diego Animal Care and Use Committee. Sprague-Dawley strain wild-type rats (Envigo/Harlan) were used for primary neuron cultures. Timed pregnant female rats (embryonic days 13-16) were singly housed in a room with a 12-hour light/dark cycle with *ad libitum* food, water access, and environmental enrichment.

### Respirometry measurements with primary cortical neurons

Cortical neurons were isolated from rat embryos (E18) as previously described ^13^. Oxygen consumption rates were measured using an Agilent Seahorse XFe96 Analyzer and Seahorse XF Mito Stress Test protocol, from 12-15 days in vitro (DIV) primary cortical neurons. Neurons were cultured on XF96e plates at 45,000/well density. Before plating, plates were coated with 20 µg/mL Poly-L-Lysine (Sigma-Aldrich) and 3.5 µg/mL Laminin (Life Technologies) overnight at room temperature. Primary neuron cultures were maintained in Neurobasal medium containing 5mM glucose, supplemented with B27, GlutaMAX, and penicillin/streptomycin (Life Technologies). Each independent experiment was performed by the preparation of new primary neuron cultures. Cortical neurons were treated with 1 µM Ara-C (Cytarabine, Tocris) at 3 DIV to prevent glia proliferation for two days. Ara-C-containing medium was replaced with fresh neuron maintenance medium at 5 DIV. The primary neuron cultures were maintained for 12-15 days in vitro by replacing one-third of the culture medium with a fresh medium every three days^14,15^. For respirometry measurements, the neuronal maintenance medium was exchanged with XF DMEM Base Medium (pH 7.4) with no phenol red (Agilent) supplemented with 5 mM glucose and 1 mM pyruvate. Respiration was measured under basal conditions as well as after injections of 1 µM Oligomycin (Sigma-Aldrich), 0.5 µM FCCP (Sigma-Aldrich), and 0.5 µM Rotenone (Sigma-Aldrich)/0.5 µM Antimycin A (Sigma-Aldrich) injections. Basal and maximal respiration values were calculated as previously described ^16^ after cell count normalization with FluxNorm. After each assay, the neuronal enrichment percentage (was calculated by immunostaining of seahorse plates with neuron-specific antibody (anti-NeuN) and cellular nuclei (NucBlue). Based on our analysis, neurons represent 60–80% of the cellular population in our cultures.

### Respirometry measurements with cell lines

C2C12 and HEK293T cells were obtained from the American Type Culture Collection (Manassas, VA) and cultured in Dulbecco’s Modification of Eagle’s Medium (DMEM) supplemented with 10% heat inactivated fetal bovine serum (Sigma-Aldrich), 1% penicillin-streptomycin and 2mM glutamine (Thermo Fisher Scientific). Oxygen consumption rates were measured using an Agilent Seahorse XFe96 Analyzer and Seahorse XF Mito Stress Test protocol. Cells were cultured 24 hours on XF96e plates at densities ranging from 2,000-20,000 cells/well for C2C12 cells, and 5,000-30,000 cells/well for HEK293T cells. For respirometry measurements, the cell maintenance medium was exchanged with XF DMEM Base Medium (pH 7.4) with no phenol red (Agilent) supplemented with 1mM sodium pyruvate (Corning Cellgro), and 1mM L-glutamine (Thermo Fisher Scientific). Respiration was measured under basal conditions as well as after injections of 1µM oligomycin, 2µM FCCP, 0.5µM rotenone/antimycin A. Basal and maximal respiration values were calculated as previously described ^16^ after cell count normalization with FluxNorm.

### Nuclei staining and imaging

Nuclei staining of cortical neurons, C2C12 and HEK293T cell were performed with NucBlue™ Live ReadyProbes Reagent (Hoechst 33342) (Invitrogen) according to the protocol provided by the manufacturer. For labeling live-cell, Hoechst 33342 dye added to XF DMEM Base Medium during the media exchange step for 15 min. For fixed cell staining, cells were first fixed with 4% paraformaldehyde for 10 minutes at room temperature, then washed three times with PBS and stained with Hoechst 33342 dye. Images were acquired using a EVOS Cell Imaging System microscope immediately after the respirometry measurements.

### FluxNorm Implementation

#### FluxNorm Overview

FluxNorm is a freeware-based method for normalization of metabolic flux data. The pipeline consists of collecting a few sample images from a well, counting the cells in those images, then extrapolating those counts to represent the entirety of the well. This methodology saves time by removing the need to take images and count cells from the entire well, while retaining accurate cell count estimates with which to normalize data. Imaging can be performed on any fluorescent microscope with sufficient resolution to distinguish cell nuclei from each other. Cell counting is performed semi-automatedly by an ImageJ macro, NucJ, which thresholds the image and counts all particles larger than a specified size. The user must define the thresholding algorithm and the minimum cell size cutoff to obtain accurate counts. Counts obtained from NucJ are then fed into a python script, ExtraPy, to be extrapolated to represent the whole well. This is done by taking the area imaged and scaling it to the seeding area of the well, ensuring counts are independent of the size of the area imaged. Final counts from ExtraPy can then be used to normalize metabolic flux data.

#### Thresholding Determination/Binarization of Images

Thresholding is an important step to acquiring accurate counts. The thresholding algorithm determines what components of the image are counted. Too low of a threshold will result in the inclusion of excess noise, while too high of a threshold will exclude actual cells. An image will need to have the resolution and signal-to-noise ratio sufficient to distinguish cells from background signal. ImageJ has a large number of built in thresholding algorithms. NucJ allows the user to select from two of these thresholding algorithms, Otsu and Huang, which are ideal for nuclei segmentation. Otsu is a great default thresholding algorithm for cell segmentation as it tries to maximize the separability of two classes in the image. This results in optimizing for separating the population of high intensity cells from the low intensity background. When the signal-to-noise ratio is high the Otsu algorithm very accurately pulls out cells from the background. The other algorithm, Huang, is more ideal when the signal-to-noise ratio is lower. Huang attempts to minimize the fuzziness of the image, which consists of trying to categorize each pixel based on its directly neighboring pixels. This results in the Huang algorithm pulling out more signal from the image compared to an algorithm like Otsu, making it ideal for situations where signal is fainter. Lastly, NucJ also enables the user to manually threshold images if neither algorithm provides adequate results. The user can adjust the algorithm of each image by hand, though this may be cumbersome if dealing with a high volume of input images. This also gives the user access to all the other thresholding algorithms built into ImageJ.

#### Cell Size Cutoff Calibration

Filtering of noisy particles was achieved by setting a cutoff that excludes cells below a minimum size. This cell size cutoff (CSC) allows the user to tune the sensitivity of NucJ to different cell types and imaging conditions. To determine an optimal CSC, we ran NucJ on preliminary data at a range of CSCs and then compared those counts to manually obtained cell counts. We then ran a python script, CaliPy, on our manually obtained and NucJ generated counts to determine the CSC that best fit our imaging conditions. CaliPy achieves this by running an ANOVA on the input data and selecting the NucJ generated counts that differ the least from the manually obtained counts.

#### Extrapolation Methodology

To avoid the time-consuming task of imaging each entire well of a seahorse plate, we opted to instead sample representative areas of the well and then extrapolate that data to represent the entire well for normalization. The formula we used for this extrapolation is:

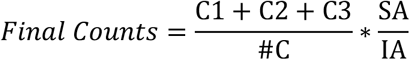

We sampled three areas (#C) from the center of each seeded well, as the metabolic flux analysis is based on the response from these cells. These areas were counted in NucJ (C1, C2, and C3) and then averaged to compute an average count for the well. Then, we scaled the count by the ratio of the Seeding Area (SA) and Imaging Area (IA). This results in an estimated count for the well which captures the most representative portion of the well.

## Data and Code Availability

FluxNorm user manual and code is available at https://github.com/pekkurnazlab/FluxNormalyzer. Any additional information required to reanalyze the data reported in this paper is available from the lead contact upon request.

## Acknowledgements

We gratefully acknowledge the invaluable contributions of the Pekkurnaz laboratory members, especially Dr. Zachary Whiddon. We also extend our appreciation to Dr. Orian Shirihai’s laboratory for their valuable insight. This project was made possible by the support of a grant from the National Institutes of Health (NIH) to G.P. (R35GM128823).

